# Dual BRD4 and AURKA Inhibition is Synergistic against MYCN-amplified and nonamplified Neuroblastoma

**DOI:** 10.1101/276840

**Authors:** Joshua Felgenhauer, Laura Tomino, Julia Selich-Anderson, Emily Bopp, Nilay Shah

## Abstract

A majority of cases of high-risk neuroblastoma, an embryonal childhood cancer, are driven by MYC or MYCN-driven oncogenic signaling. While considered to be directly “undruggable” therapeutically, MYC and MYCN can be repressed transcriptionally by inhibition of Bromodomain-containing protein 4 (BRD4) or destabilized posttranslationally by inhibition of Aurora Kinase A (AURKA). Preclinical and early-phase clinical studies of BRD4 and AURKA inhibitors, however, show limited efficacy against neuroblastoma when used alone. We report our studies on the concomitant use of the BRD4 inhibitor IBET-151 and AURKA inhibitor alisertib. We show that, in vitro, the drugs act synergistically to inhibit viability in three models of high-risk neuroblastoma. We demonstrate that this synergy is driven, in part, by the ability of IBET-151 to mitigate reflexive upregulation of AURKA, MYC, and MYCN in response to AURKA inhibition. We then demonstrate that IBET-151 and alisertib are effective in prolonging survival in three xenograft neuroblastoma models in vivo, and this efficacy is augmented by the addition of the antitubule chemotherapeutic vincristine. These data suggest that epigenetic and posttranslational inhibition of MYC/MYCN-driven pathways may have significant clinical efficacy against neuroblastoma.

**Abbreviations:** AURKA, aurora kinase A; BRD4, Bromodomain-containing protein 4; CI, combination index; F_a_, fraction affected; LOH, loss of heterozygosity; 11q, long arm of chromosome 11

**Funding sources:** This work was supported by the Families for a Cure foundation (grant number 20064014) and CancerFree Kids (grant number 82178416).

## INTRODUCTION

Advanced neuroblastoma, the embryonal childhood cancer arising from sympathoadrenal precursors, remains a major clinical challenge. Patients with high-risk tumors at diagnosis are treated with aggressive multimodal chemotherapies, radiation therapy, and immunotherapy but suffer high rates of disease progression and/or recurrence, with cure rates ∼50%^1,2^. Those patients with progressive neuroblastoma rarely have durable responses to current salvage therapies and die of disease^3^. *MYCN* and/or *MYC* amplification or overexpression have been shown to be oncogenic drivers in a majority of advanced neuroblastomas^4-7^. These proteins are transcription factors, promoting expression of numerous oncogenes and enhancing cell proliferation and survival^8^, but also function as repressors of cell signaling^9,10^ and as drivers of transcriptional elongation and activation of superenhancers through interactions with CDK7/9 and RNA Polymerase II^11-13^.

MYC and MYCN are difficult to therapeutically target directly, but novel agents have been designed to destabilize or repress these oncoproteins indirectly. One class of drugs against MYC/MYCN-driven cancers target Aurora Kinase A (AURKA), a protein with multiple functions in cytokinesis ^14^ and in the stabilization of MYC and MYCN by prevention of FBXW7-mediated ubiquitination^15^. The first-in-class drug, alisertib, showed efficacy against neuroblastoma, particularly MYCN-amplified disease, preclinically^16^. However, in the Phase 1 pediatric clinical trial, it had higher toxicity in children than in adults, limiting its maximally tolerated dose^17^. Alisertib failed to meet response criteria in multiple phase 2 studies when used alone^18-21^ but is being examined in combination therapies.

A second class of drugs against MYC/MYCN-driven cancers inhibits the bromodomain and extraterminal motif (BET) chromatin-binding proteins. These proteins recognize and localize to acetylated lysine residues^22^ and promote transcription by recruiting and phosphorylating components of RNA Polymerase II^23^. One BET protein, BRD4, has been shown to be active in cancers by promoting expression of multiple targets, including *CDK4/6*^24^, *BCL2*^25^, *MCL1^26,27^, MYC*^28^ and *MYCN*^29^. BRD4 inhibitors, developed for research and clinical use, have shown some preclinical efficacy against *MYCN-*amplified neuroblastoma but did not induce regression when used alone^25,29,30^. The cytostatic effects of BRD4 inhibitors suggest that these drugs may have limited effects clinically when used alone, particularly in diseases where BRD4 supports oncogenesis but is not the primary disease driver.

AURKA and BRD4 inhibitors both attack many common oncogenic drivers in distinct but complementary ways. In this study, we show that the AURKA inhibitor alisertib and the BRD4 inhibitor IBET-151 have significant synergy against neuroblastoma cell lines in vitro, inhibiting viability at significantly lower doses than when either drug is used alone. We show that cells treated with alisertib have a reflexive transcriptional upregulation of *AURKA, MYC*, and *MYCN*, but concomitant treatment with IBET-151 represses that upregulation. Treatment with both drugs is more effective at repressing expression of multiple oncoproteins, including MYC, MYCN, CDK4/6, AURKA, and BCL2. In three tumor xenograft models, IBET-151 and alisertib are more efficacious together in extending survival than either drug alone and induce tumor regression in a *MYCN*-amplified model. Furthermore, the addition of the anti-tubulin chemotherapeutic vincristine augments this effect, inducing durable tumor regression that is maintained after cessation of treatment in a *MYCN*-amplified model and a *MYCN*-nonamplified model and extending survival in a third *MYCN*-nonamplified model.

## MATERIALS AND METHODS

### Cell Lines

SK-N-SH cell line was obtained from Javed Khan (National Cancer Institute, Bethesda, MD). SK-N-AS cell line was obtained from American Type Culture Collection (Manassas, VA). NB1643 and NB-SD cell lines were obtained from Peter Houghton (Greehey Children’s Cancer Research Institute, San Antonio, TX). All cell lines were authenticated by PowerPlex16 short tandem repeat analysis (Promega) at the start of in vitro studies and again prior to in vivo studies. Cells were cultured in DMEM (Corning, Bedford, MA) with 10% FBS (PeakSerum, Wellington, CO) at 37°C with 5% CO_2_ and confirmed to be free of *Mycoplasma* by SouthernBiotech Mycoplasma Detection Kit (Birmingham, AL), tested every 3 months.

### Drugs

Alisertib was purchased from ApexBio (Houston, TX). IBET-151 was obtained from GlaxoSmithKline (Collegeville, PA). A list of primers and antibodies used can be found in the supplementary data.

### Cell viability assay and Combination Index (CI) analysis

NB-1643, SK-N-SH, and SK-N-AS cells were plated in 96-well plates at 25,000, 25,000, and 5,000 cells/well respectively in complete media in triplicate wells for each dose and cultured for 24 hours. Cells were treated with either IBET-151 dissolved in DMSO with concentration from 20-2000 nM, alisertib dissolved in ethanol with concentrations from 10-1000 nM), both drugs, or vehicle control for 48 hours. Cell viability was measured using the IncuCyte ZOOM® live cell imaging system (Essen BioScience, Ann Arbor, MI) to track percent confluence of each well. Percentage confluence as compared to vehicle control was used to calculate treatment effect. IC50 and combination index (CI) values were calculated using Compusyn software (Combosyn, Inc., Paramus, NJ). Three independent experiments were performed; representative experiments are shown here.

### Reverse Transcription – quantitative Polymerase Chain Reaction (RT-qPCR)

Cells were grown to 80% confluence, then treated with 1 mcM IBET-151, 1 mcM alisertib, both drugs at 1 mcM, or vehicle control for 24 hours. Total RNA was extracted from the cells using NucleoSpin RNA purification kit (Takara Bio USA), and 1 mcg of RNA used for cDNA synthesis using Maxima RT cDNA First Strand Synthesis kit (ThermoFisher Scientific, Waltham MA). qPCR was performed using KiCqStart SYBR Green qPCR ReadyMix (Sigma-Aldrich, St. Louis, MO) using the ABI PRISM 7900HT thermal cycler (ThermoFisher Scientific), with relative quantitation by the ddC_t_ method as previously described^31^. Experiments were performed with technical duplicates on each plate and in three independent experiments, with the relative expression of each experiment used to calculate expression and standard deviation, plotted on each graph.

### Western blot

Cells were grown to 80% confluence, then treated with 1 mcM IBET-151, 1 mcM alisertib, both drugs at 1 mcM, or vehicle control for 48 hours. Cells were collected by scraping and lysed using RIPA buffer, with 50 mcg of protein/sample used for western blot as previously described^31^. Blots were imaged by chemiluminescence using ECL Western Blotting Substrate (ThermoFisher Scientific). Experiments were performed in independent triplicate; representative images are shown here. Complete blots are shown in the Supplementary Data.

### Tumor xenograft studies

5×10^6^ cells of each type were suspended in PBS and mixed 1:1 in Matrigel (Corning) to a final volume of 100 mcL and injected subcutaneously into the flanks of SCID mice (Envigo, Indianapolis, IN), one injection per mouse. Tumors were grown to ∼200 mm^3^ as estimated by volume=(length x width^2^)/2. Mice were then treated with the listed drug combinations, n=5 per group, with drugs at the following doses and routes: IBET-151, injected intraperitoneally 20 mg/kg/day; alisertib, orally by gavage 20 mg/kg/day; vincristine, injected intraperitoneally 0.1mg/kg/dose once weekly (formulations in the Supplementary Data). Mice were treated 5 days x5 weeks, then observed without treatment for 2 weeks. Mice were weighed and tumors measured twice weekly. Mice were euthanized at humane endpoints, when tumors attained 2000 mm^3^, or at the end of the study, with tumors harvested. All studies were designed in accordance with Nationwide Children’s Hospital Institutional Animal Care and Use Committee (IACUC) guidelines and performed under IACUC-approved protocols.

### Statistical analysis

All statistical analyses were completed using GraphPad Prism 7 (GraphPad Software, Inc., La Jolla, CA). Where appropriate, the two-tailed Student’s t-test was used to calculate significant differences between comparison groups in the experiments above. For multiple comparisons, one-way analysis of variance (ANOVA) was used with the Bonferroni correction for multiple comparisons against a single control. Mantel-Cox log rank test used for survival analyses.

## RESULTS

### IBET-151 and Alisertib Synergistically Kill Neuroblastoma Cells In Vitro

We first aimed to demonstrate if IBET-151 and alisertib had synergistic antineoplastic activity in vitro. We used SK-N-AS and SK-N-SH, *MYCN*-nonamplified cell lines, and NB1643, a *MYCN-*amplified cell line. We treated the cells with a range of doses of IBET-151 from 0-2000 nM and with alisertib from 0-1000 nM for 48 hrs, defining IC50 doses for each drug and cell line (Supplementary Table 1). All cell lines were sensitive to both drugs, although alisertib was much more cytotoxic at 48 hours than IBET-151, which appeared to be cytostatic morphologically. We then treated the cells with all combinations of those doses and evaluated cell viability at 48 hrs. Using Compusyn software, we found that, in all three cell lines, most drug combinations used were significantly synergistic (Figure 1). The Compusyn software calculates, for each drug dose combination, the expected effects of the drugs alone and compares this to the observed effect, calculating a combination index (CI)^32^. CI<1 shows a greater-than-additive effect of the drugs, and CI<0.7 is considered synergistic. For all three cell lines, most drug dose combinations had CI<0.7 and as low as 0.1. This supported our initial hypothesis that combined BRD4 and AURKA inhibition synergistically inhibit neuroblastoma viability.

**Figure 1:**
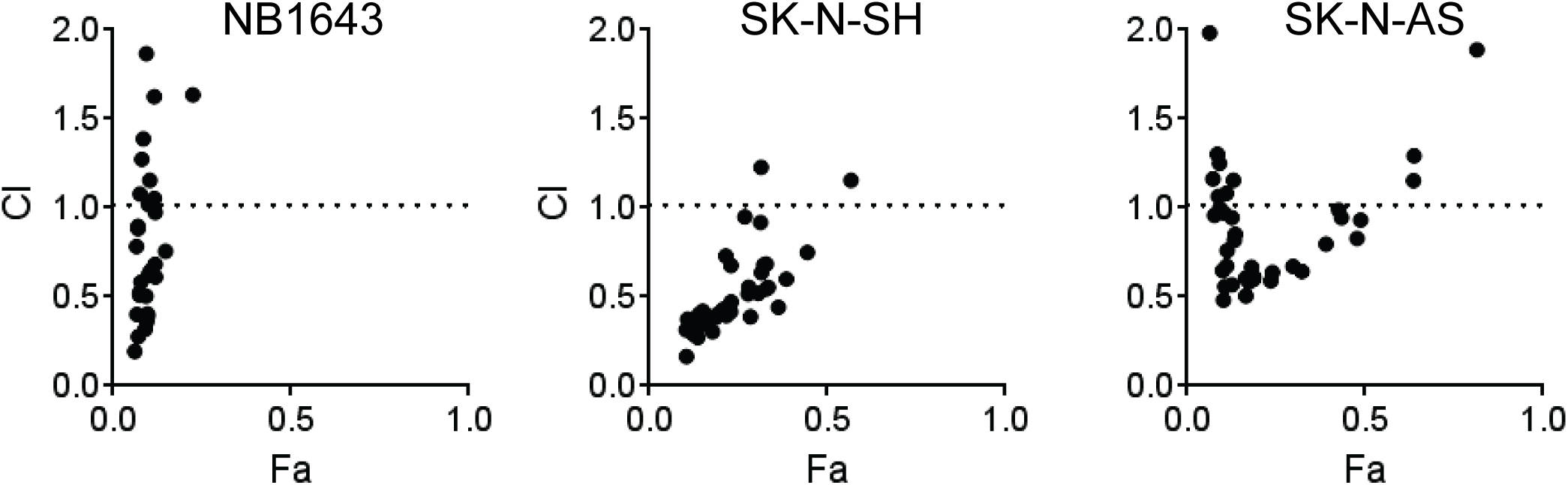
IBET-151 and alisertib are synergistic in their effects on neuroblastoma cell viability in vitro. Combination Index (CI) plots are shown for NB1643, SK-N-SH, and SK-N-AS neuroblastoma cell lines. Cells were treated with a combination of doses of IBET-151 and alisertib for 48 hours, as described in the methods, and images were obtained to quantify percentage confluence per well as a proxy for cell viability, here quantified as the F_a._ (x-axis). Each dose combination and the resultant F_a_, as well as the F_a_ for each drug alone at individual doses, were entered into the Compusyn software. From these data, the Combination Index (CI) for each dose combination was calculated, determined if the F_a_ observed was antagonistic (less than each drug’s effect alone, CI>1), additive (equal to the effective of each drug alone, CI=1), or greater-than additive (greater than the effect of each drug alone, CI<1). CI<0.7 is generally considered synergistic. For each cell line, a majority of drug combinations had CI<1, with F_a_<0.5 for all such combinations. Representative experiments shown.

### IBET-151 and Alisertib Have a Greater Effect in Combination in Repressing Target Protein Expression

We hypothesized that IBET-151 and alisertib exerted synergy due to complementary mechanisms of activity, namely that IBET-151 would repress transcriptional expression of those proteins posttranslationally repressed by alisertib. We examined the transcriptional expression of a number of the gene targets of these drugs. SK-N-AS, SK-N-SH, and NB1643 cells were treated with 1 mcM of IBET-151, alisertib, both, or equal volume of vehicle, for 24 hours, then harvested for RNA used for RT-qPCR. For most genes tested, we observed that treatment of the cells with IBET-151 repressed gene expression at 24 hours below levels seen in controls, whereas treatment with alisertib alone resulted in upregulation of those target genes, particularly *AURKA, MYC*, and *MYCN* (Figure 2). Use of IBET-151 with alisertib mitigated that upregulation, preventing compensation for AURKA inhibition (one-way ANOVA for SK-N-AS among the 4 treatment groups p=0.0079, for SK-N-SH p=0.0482, for NB1643 p=0.0169). The degree of these effects varied among the cell lines. In the *MYCN-*amplified NB1643 cells, IBET-151 had a modest effect on repression of gene expression alone, alisertib caused prominent rebound gene expression, but IBET-151 with alisertib had limited rebound gene expression for most genes tested, except for *AURKA* itself. The *MYCN*-nonamplified SK-N-SH cells had similar patterns of expression in response to the drugs, though with less rebound gene expression in response to alisertib. The *MYCN*-nonamplified SK-N-AS cells had the least amount of rebound gene expression in response to alisertib but had more consistent repression of gene expression by IBET-151, alone or with alisertib.

**Figure 2:**
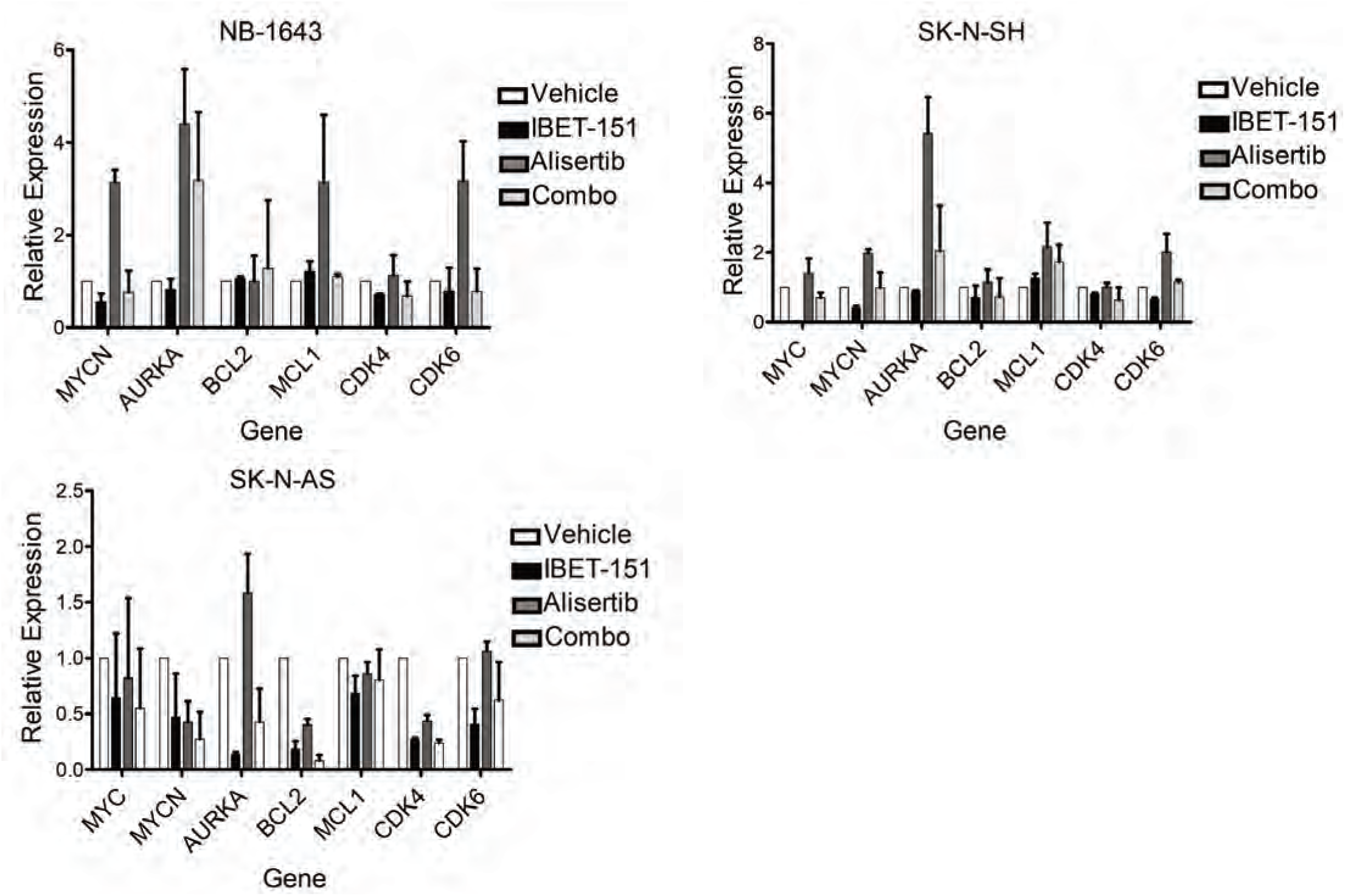
AURKA inhibition with alisertib induces increases RNA expression of oncogenes, which is mitigated by BRD4 inhibition with IBET-151. Cells were treated with 1 mcM IBET-151, 1 mcM alisertib, both drugs, or vehicle control for 24 hours, after which RNA was extracted from the cells and used for RT-qPCR. Treatment with alisertib alone (dark grey bars) induced overexpression of most oncogenic targets tested in the NB1643 and SK-N-SH cells, with less of an effect on SK-N-AS cells except for *AURKA.* Co-treatment of the cells with IBET-151 reduced or entirely abrogated this overexpression for most target genes in all three cell lines. Relative expression shown using ddC_t_ methods, with each gene’s expression normalized first to housekeeping genes in each sample then against each gene’s expression in the vehicle control. Three independent experiments performed, with mean and standard error plotted, one-way ANOVA based on treatment for SK-N-AS among the 4 treatment groups p=0.0079, for SK-N-SH p=0.0482, for NB1643 p=0.0169.

We confirmed the effects of this drug combination on protein expression by western blot. In all three cell lines, we found that IBET-151 with alisertib was more effective than alisertib alone in repressing expression of multiple oncoproteins, including AURKA, MYC, MYCN, and CDK4/6 (Figure 3), mitigating reflexive protein overexpression in reaction to alisertib. More variable effects were seen on BCL2 and MCL1, with MCL1 repressed in NB1643 and SK-N-SH cells, BCL2 additionally repressed in SK-N-SH cells, but neither MCL1 nor BCL2 significantly affected by IBET-151 and/or alisertib in SK-N-AS cells. These data further supported our hypothesis that the two drugs can synergistically repress expression of common oncoprotein targets.

**Figure 3:**
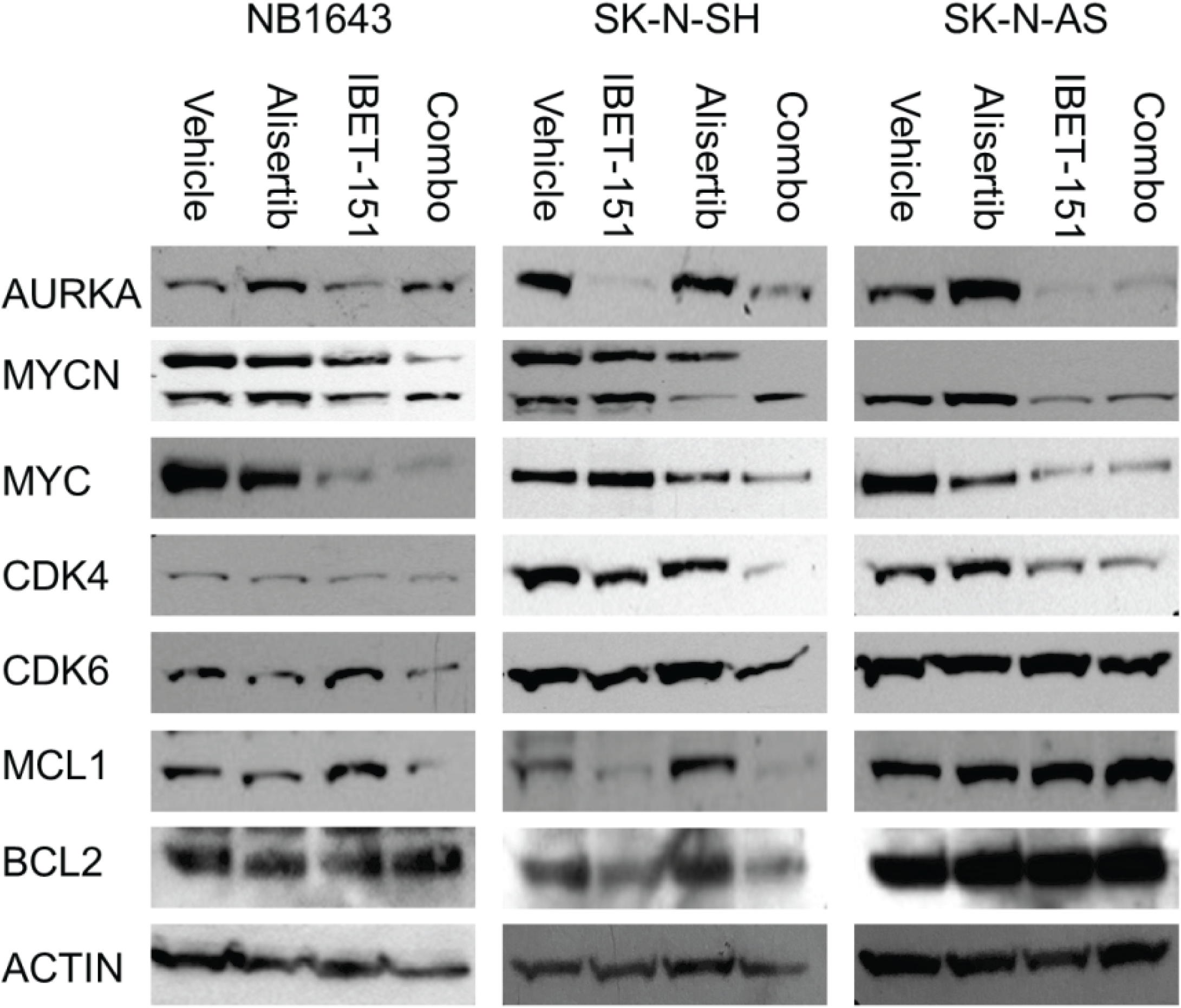
Dual AURKA and BRD4 inhibition is most efficacious in repressing target oncoprotein expression. Western blots of oncoprotein expression in three neuroblastoma cell lines. Cells were treated with 1 mcM IBET-151, 1 mcM alisertib, both drugs, or vehicle alone for 48 hours, after which cells were lysed and total protein harvested for western blot. For all three cell lines, treatment with the AURKA inhibitor alisertib caused increased total AURKA expression, but with appreciable decreases in MYC and/or MYCN expression. Treatment with BRD4 inhibitor IBET-151 caused more obvious decrease in AURKA expression, but with more variable effects on MYC/MYCN expression. Use of both inhibitors caused greater decrease in protein expression of AURKA, MYCN, and MYC as compared to alisertib alone in all three cell lines. Similar changes were seen on CDK4/6 and MCL1 in NB1643 and SK-N-SH cells, but SK-N-AS cells showed no obvious effects on CDK6, MCL1 or BCL2 expression with any drug treatments. Three individual experiments were performed, with representative blots shown.

### IBET-151 and Alisertib have greater efficacy together against neuroblastoma tumor xenografts in vivo than either drug alone

Given these preliminary data, we next evaluated the efficacy of these drugs in vivo. SK-N-SH, NB1643, and SK-N-AS cells were implanted subcutaneously into the flanks of SCID mice to generate tumor xenografts. When the xenografts were 200 mm^3^ in volume, the mice were treated with either IBET-151, alisertib, both drugs together, or vehicle alone, to humane endpoint or for 5 weeks, with surviving mice observed for a drug washout period of 2 weeks. Each group of mice tolerated the treatments well, with no significant weight loss or indications of physiologic stress.

In the mice with SK-N-SH tumors, treatment with IBET-151 did not extend survival as compared to vehicle (Figure 4A, median survival 38 days vs 38 days, p=0.47). Treatment with alisertib did extend survival (median survival 56 days, p=0.0064), but only one mouse survived throughout the study. IBET-151 and alisertib together were superior to either drug alone (medial survival undefined, p=0.0018 vs IBET-151, p=0.0416 vs alisertib), with 4/5 mice surviving through the washout period, though their tumors did regrow after treatment ended (Figure 5A).

**Figure 4:**
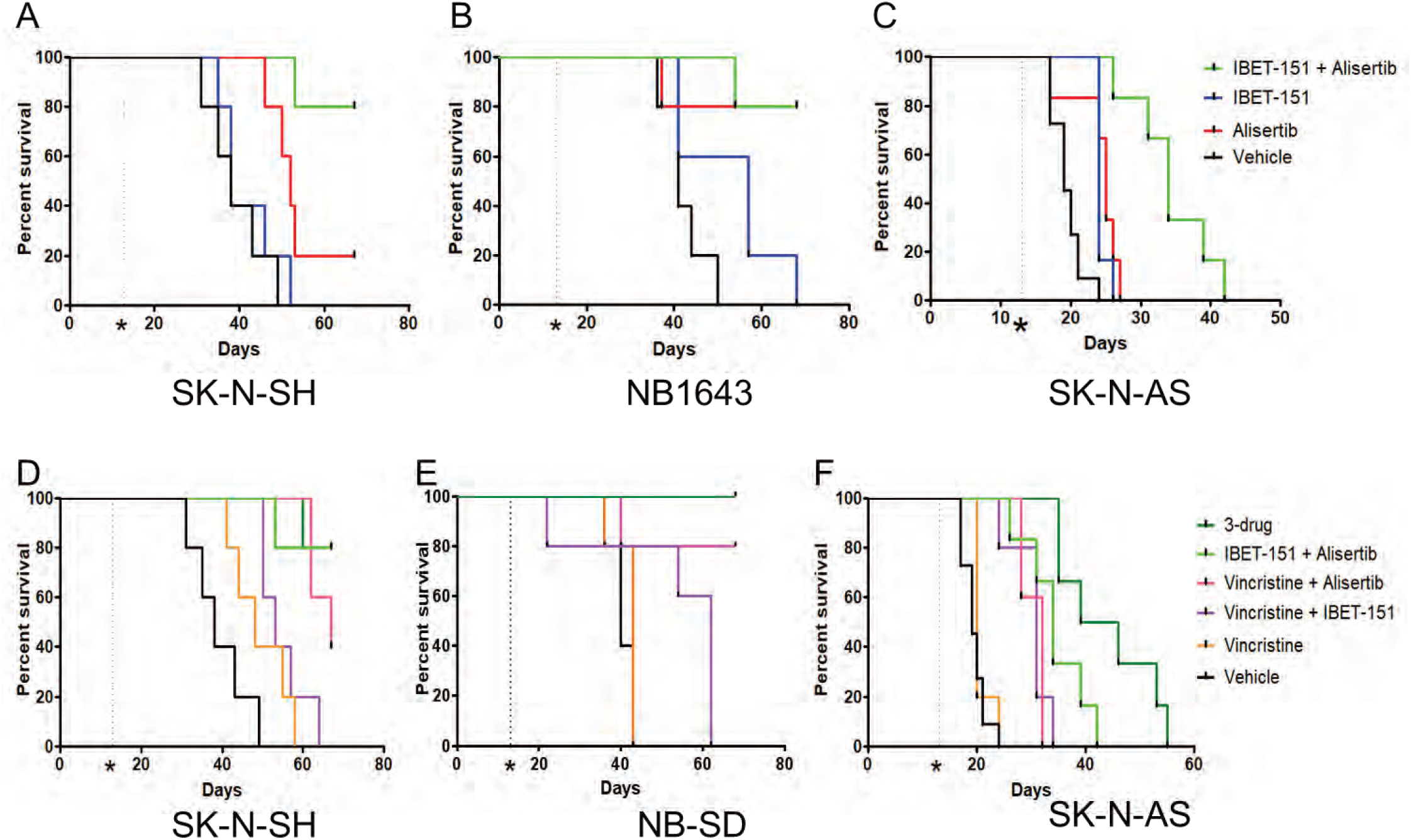
Drug combinations of IBET-151 and alisertib, with or without vincristine, improve survival of mice with neuroblastoma tumor xenografts. Kaplan-Meier survival curves of tumor xenograft studies. See main text for description of treatment methods, n=5 mice per treatment group. Dashed vertical line with asterix shows timepoint after xenograft injection at which treatment was started for all groups. For mice treated with IBET-151, alisertib, both drugs, or vehicle alone, the two-drug combination was most efficacious as compared to vehicle control in extending survival (see main text for individual comparisons and p-values, calculated by Mantel-Cox logrank test). For mice treated with various combinations of IBET-151, alisertib, vincristine and vehicle, the three-drug combination was most efficacious as compared to vehicle in extending survival (see main text for individual comparisons and p-values, calculated by Mantel-Cox logrank test).

**Figure 5:**
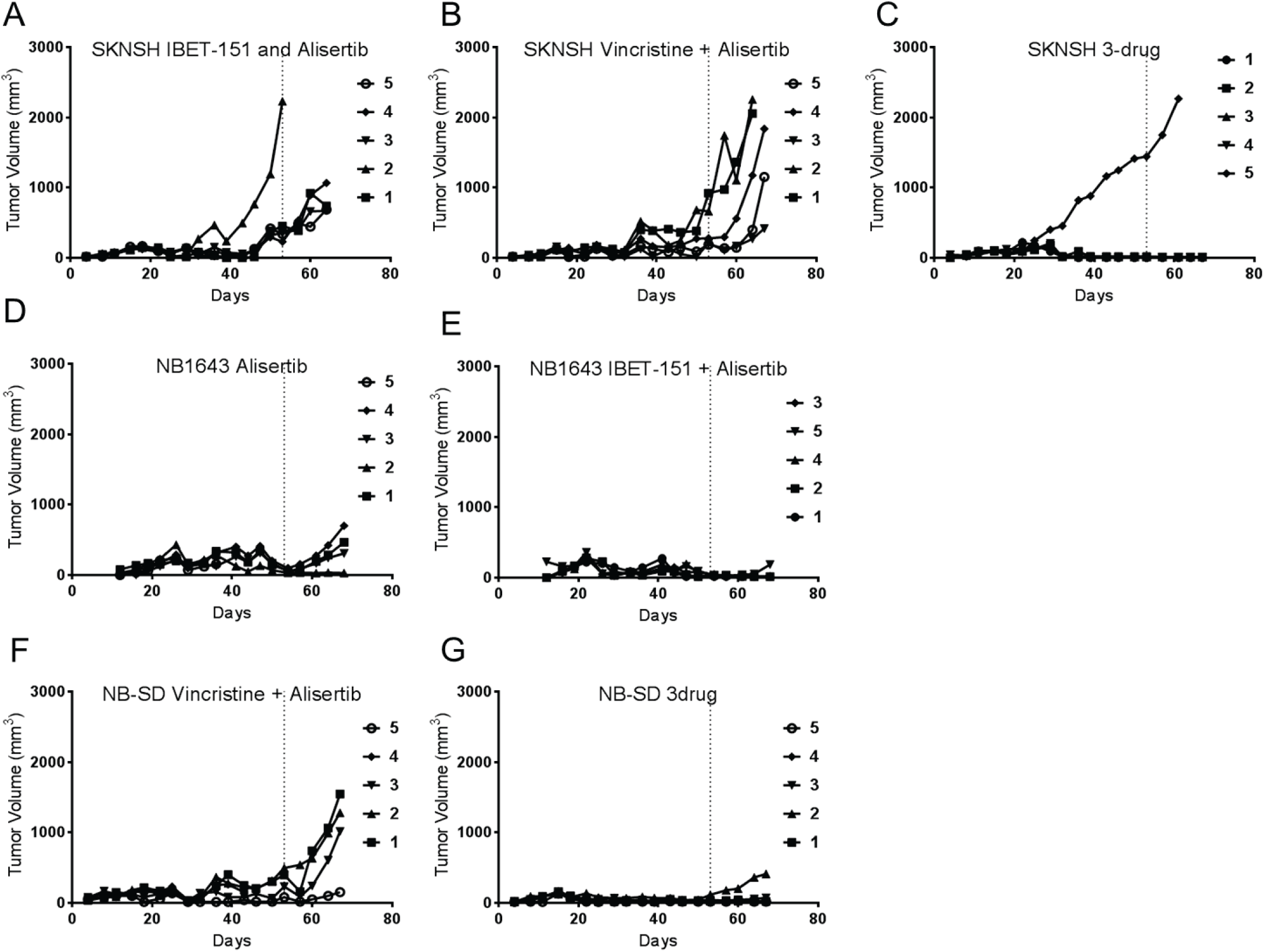
The combination of IBET-151, alisertib, with or without vincristine was most efficacious in inducing durable tumor regressions in neuroblastoma tumor xenografts in vivo. Tumor volume plots of individual tumor xenografts throughout the experiments. In SK-N-SH xenografts, treatment with alisertib and either IBET-151 (A) or vincristine (B) extended survival but tumors grew despite treatment, particularly after treatments ended and mice were monitored (vertical dashed line), but treatment with all three drugs maintained durable tumor regression even after treatment ended (C). In NB1643 xenografts, alisertib alone was efficacious in slowing tumor growth and extending survival, but tumors grew nonetheless (D), whereas treatment with IBET-151 and alisertib was more efficacious in maintaining durable regression (E). Similarly, in NB-SD xenografts, treatment with vincristine and alisertib slowed tumor growth but did not induce durable tumor regression (F), but treatment with all three drugs did (G).

In the mice with NB1643 tumors, treatment with IBET-151 trended toward significantly extending survival as compared to vehicle alone (Figure 4B, median survival 57 days vs 41 days, p=0.069). Treatment with alisertib strongly significantly extended survival as compared to vehicle (median survival undefined, p=0.0090), with 4/5 mice surviving through therapy and washout. Treatment with IBET-151 and alisertib similarly extended survival as compared to vehicle (median survival undefined, p=0.0018). However, tumor growth was different between those mice treated with alisertib alone as compared to with both drugs (Figure 5D and E). Of the mice treated with alisertib alone, 3/4 mice had their tumors regrow after the end of treatment. In contrast, only 1/4 mice treated with both drugs had its tumor grow appreciably after treatment ended, suggesting the drug combination had a more durable antitumor effect against this MYCN-amplified tumor xenograft model.

In the mice with SK-N-AS tumors, treatment with either IBET-151 or alisertib significantly extended survival as compared to mice treated with vehicle (Figure 4C, median survival 24 vs 19 days, p=0.0009, for IBET-151 vs vehicle, and median survival 25 vs 19 days, p=0.0038, for alisertib vs vehicle). Treatment with both drugs together was superior to treatment with either drug alone (median survival 34 days, p<0.005 as compared with vehicle or either drug alone), though the combination did not halt tumor growth fully in any of these mice. Nonetheless, these data were supportive of improved efficacy of BRD4 and AURKA inhibition in combination against all subtypes of neuroblastoma.

### Addition of Vincristine to IBET-151 and Alisertib improves Antitumor Efficacy and Induces Durable Complete Responses in Neuroblastoma Tumor Xenografts

The results of the tumor xenograft studies suggested that there was either a delayed or incomplete cytotoxic effect of the drug combination against neuroblastoma in vivo, but that addition of a cytotoxic chemotherapeutic could improve the efficacy of the drug combination. Vincristine, a vinca alkaloid that inhibits tubulin polymerization, has been shown to be synergistic in other cancers with both BRD4 inhibitors^33^ and AURKA inhibitors^34^. For this and additional reasons discussed below, we evaluated the efficacy of vincristine with IBET-151, with alisertib, and in combination with both drugs.

A pilot study identified that, while mice tolerated vincristine dosing of 0.2 mg/kg intraperitoneally weekly alone or with alisertib, they did not tolerate that dose in combination with IBET-151, with increased vincristine toxicity, urinary retention and weight loss. Accordingly, we reduced the vincristine dosage to 0.1 mg/kg weekly. Given the high sensitivity of NB-1643 xenografts to alisertib alone and with IBET-151, we instead used xenografts derived from the NB-SD cell line, another MYCN-amplified cell line with reported greater resistance to alisertib^35^.

Mice harboring SK-N-SH xenografts had a modest benefit from treatment with vincristine alone vs vehicle control (median survival 48 days vs 38, p=0.08, Figure 4D). Vincristine with IBET-151 did significantly extend survival vs vehicle (median survival 53 days, p=0.0018) but not more than vincristine alone (p=0.33). Vincristine with alisertib extended survival as compared to vincristine alone or with IBET-151 (median survival 67 days, p=0.0018 vs vehicle or vincristine, p=0.015 vs vincristine+IBET-151), though not significantly better that IBET-151 with alisertib (p=0.32). Vincristine with alisertib did maintain survival throughout the treatment period for all mice, but all tumors regrew during the washout period (Figure 5B). Treatment with all three drugs was highly efficacious at extending survival as compared to vincristine with or without IBET-151 (median survival undefined, p=0.0018 vs vincristine or control, p=0.0062 vs vincristine +IBET-151) though not significantly better than alisertib with IBET-151 (p=0.9372) or with vincristine (p=0.32). However, in striking contrast to all other treatment, use of vincristine, alisertib and IBET-151 maintained repression of tumor growth through the washout period for 4/5 mice, inducing durable complete regressions (Figure 5C).

Mice harboring NB-SD xenografts responded similarly to the drug combinations. Vincristine did not significantly improve survival compared to vehicle (median survival 43 vs 40 days, p=0.2845, Figure 4E). Vincristine with IBET-151 improved survival vs vehicle (median survival 62 days, p=0.044) and trended toward significant improvement vs vincristine alone (p=0.059). Vincristine with alisertib significantly improved survival compared to these treatments (median survival undefined, p=0.02 vs vehicle, p=0.046 vs vincristine, and p=0.041 vs vincristine and IBET-151). Four of the five mice treated with vincristine and alisertib survived the entire study, but all mice had tumor growth during the washout period (Figure 5F). The three drugs together further improved survival (median survival undefined, p=0.0026 vs vehicle, p=0.0035 vs vincristine, p=0.0034 vs vincristine and IBET-151, p=0.32 vs vincristine and alisertib). More importantly, all 5 mice survived the entire study, 4 of 5 mice had no significant regrowth during the washout period, and only one tumor had growth during the washout period but still remained <500 mm^3^ (Figure 5G).

Mice harboring SK-N-AS xenografts also benefited from the addition of vincristine, though to a less pronounced degree. Vincristine did not improve survival compared to vehicle (median survival 20 vs 19 days, p=0.27). Vincristine with IBET-151 was significantly more efficacious than vehicle or vincristine (median survival 31 days, p=0.0008 vs vehicle, p=0.0047 vs vincristine). Vincristine with alisertib improved survival (median survival 32 days) as compared to vehicle (p=0.0004) or vincristine (p=0.0016) but not as compared to vincristine and IBET-151 (p=0.90), in contrast to the other xenograft models. The three-drug combination was most effective at extending survival (median survival 46 days, p=0.0004 vs control, 0.0016 vs vincristine, p=0.0020 vs vincristine and IBET-151, p=0.0031 vs vincristine and alisertib, p=0.027 vs IBET-151 and alisertib). However, while the three-drug combination slowed tumor growth, none of the mice survived through the treatment course. Nonetheless, these data support efficacy of chemotherapy with BRD4 and AURKA inhibition against multiple types of neuroblastoma.

## DISCUSSION

*MYCN* or *MYC* amplification and/or overexpression have been shown to be the oncogenic drivers in over half of high-risk neuroblastomas^4,36-38^. These transcription factors have been generally considered to be “undruggable” therapeutic targets directly, but studies into the regulation and stabilization of MYC/MYCN identified indirect approaches to impair their tumorigenic programs. Repression of transcriptional expression by BRD4 inhibition and posttranslational destabilization by AURKA inhibition both demonstrated efficacy in vitro against neuroblastoma. However, these approaches in isolation failed to have significant efficacy in clinical trials. Our data show that combined BRD4 and AURKA inhibition is synergistic in against MYC/MYCN-associated oncogenic pathways. This combination has efficacy against MYCN-amplified and nonamplified models of neuroblastoma in vivo, and this efficacy is further augmented by use of chemotherapy such as vincristine.

The comparative efficacy of BRD4 and AURKA inhibition varied in our neuroblastoma models with some association with their genomic alterations. NB1643 and NB-SD cell lines harbor *MYCN*-amplification, while SK-N-SH and SK-N-AS cell lines do not. Additionally, SK-N-AS cells have unbalanced loss of heterozygosity (LOH) of chromosome 11q, a demonstrated clinical biomarker of poor prognosis^39,40^. In the combinatorial in vitro assay, the drugs were synergistic in all three models tested but with different dose effects. In the synergy plot (Figure 1), the X-axis indicates the F_a_ value, the percentage of viable cells at the experimental endpoint. The NB1643 cells had very high sensitivity to the drugs in combination, with F_a_<0.2 for all synergistic combinations after 48 hours of treatment, with the majority of the cell death induced by alisertib. This mirrors studies that showed strong efficacy of alisertib preclinically against *MYCN*-amplified neuroblastoma^16,35^. Comparatively, the F_a_ of the drug combination in the SK-N-SH and SK-N-AS cells ranged across the synergistic doses from 0.1-0.5.

This mirrored drug efficacy in the tumor xenograft models, in which the *MYCN*-amplified NB1643 and NB-SD xenografts had significantly higher sensitivity to alisertib-including combinations as compared to SK-N-SH and SK-N-AS. We theorize that this is due to the addiction of *MYCN*-amplified neuroblastoma to the MYCN-driven oncogenic pathways, making them very sensitive to any destabilization of MYCN. There was benefit in all models with the addition of IBET-151, with the greatest impact in SK-N-SH cells, in which IBET-151-including combinations improved survival and most significantly slowed tumor regrowth. This could be because the SK-N-SH cells are less addicted to MYC/MYCN biology and more dependent on other oncogenic pathways affected by BRD4 inhibition, including cell cycle progression, cytokinesis, and anti-apoptosis, as shown in the expression analyses (Figure 2 and 3).

While the drug combination did improve survival in mic with SK-N-AS xenografts, IBET-151 and alisertib was not as efficacious in inducing tumor regression as in the other models. The expression analyses showed that there was less effect of either drug alone or together on some pathways, particularly on MCL1 expression. It is possible SK-N-AS cells may survive by activation of antiapoptotic pathways by MCL1, which may or may not be due to LOH of 11q. SK-

N-AS has also been shown to be chemotherapy-resistant^41,42^, and has particularly high expression of ABCC1, which allows for drug efflux^43^. Thus, resistance to IBET-151 and alisertib may be due to elimination of the drugs before there is sufficient exposure for durable effect. Evaluation of the resistance mechanisms against BRD4 and AURKA inhibition are warranted in other models of neuroblastoma with LOH of 11q or ABCC1 expression.

Although alisertib showed preclinical antitumor efficacy, it failed to show meaningful activity in single-agent use in multiple clinical trials ^18,44,45^, in which it was administered for 7 days with 14 days of recovery. We have shown that alisertib treatment induces rebound transcriptional overexpression of its targets, which may account for the lack of clinical efficacy. This rebound expression can be repressed by BRD4 inhibition, which we hypothesize allows for greater antitumor efficacy. This finding may impact future clinical trial design, whereby direct enzymatic inhibition, such as with alisertib, delivered in a pulsatile fashion may be best matched with chronic use of an epigenetic inhibitor, such as IBET-151, to prevent oncogenic reactivation and maintain tumor regression.

While IBET-151 and alisertib were significantly more efficacious together than alone, the combination had a delayed tumor regression effect. We hypothesized that a cytotoxic agent could rapidly induce tumor regression that could then be maintained by BRD4 and AURKA inhibition. We chose vincristine to test this hypothesis. Prior data showed the vincristine is synergistic with IBET-151 and alisertib individually^33,34^. Vincristine is used sparingly in upfront neuroblastoma therapy, due to limited single-agent efficacy, but it has improved efficacy in combination with drugs that cause cell cycle disruption^46-48^ as occurs with BRD4 and AUKRA inhibition. Furthermore, vincristine is not significantly myelosuppressive, avoiding a toxicity associated with alisertib^17^. These features suggested that vincristine could be combined with BRD4 and AURKA inhibition in clinical trials for neuroblastoma. We did not anticipate increased toxicity with combined vincristine and IBET-151, requiring vincristine dose reduction, but we were encouraged to still see increased efficacy in the three-drug combination. Identification of other chemotherapeutic agents synergistic with BRD4 and AURKA inhibition, including IBET-151, alisertib, and/or other drugs in these classes, may find other combinations for clinical translation.

This study demonstrates that combined epigenetic and posttranslational targeting of oncogenic pathways can be synergistic. Whereas posttranslational targeting may cause rapid changes in oncogenic pathways, efficacy may be limited because of the transiency of effect. Epigenetic targeting of cancer cells may allow for more global and durable effects but can be limited by the slow cytotoxic effect, allowing tumors to grow before clinical efficacy can be appreciated. Dual epigenetic and posttranslational inhibition may improve clinical efficacy and also salvage drugs that have failed primary clinical endpoints when used alone. The drugs can be used together to increase antitumor efficacy or they can also be dose-adjusted to reduce toxicity while still maintaining or improving single-agent efficacy. The specific approach to target MYC/MYCN-driven oncogenic pathways may have broader impact on a host of cancer types in which these pathways are active, including medulloblastoma^49^, rhabdomyosarcoma^50^, pancreatic neuroendocrine tumors^51^, prostatic neuroendocrine tumors^52^, and breast cancer^53^.

## CONCLUSIONS

Our results demonstrate that combined epigenetic *MYC/MYCN* inhibition by use of the BRD4 inhibitor IBET-151 and posttranslational inhibition of MYC/MYCN stabilization by use of the Aurora Kinase A inhibitor alisertib are efficacious in antitumor effects against neuroblastoma with or without MYCN amplification. This combination approach is more effective in inducing and maintaining transcriptional and protein repression of multiple oncoproteins than each drug alone, and this is a likely driver of the drug combination’s synergy. This antitumor effect is further improved with the addition of the antitubule chemotherapeutic vincristine, inducing durable tumor regressions in multiple tumor xenograft models of neuroblastoma in vivo. This study supports use of dual BRD4 and AURKA inhibition in clinical studies of neuroblastoma and other MYC/MYCN-driven malignancies.

## Acknowledgements

We thank Olena Barbash at GlaxoSmithKline for her assistance and expertise in the use of the BRD4 inhibitor IBET-151 in vitro and in vivo.

